# The potato resistance protein Rx1 multimerizes upon activation by the Coat Protein of PVX

**DOI:** 10.1101/2023.01.10.523371

**Authors:** Marijn Knip, Emy Latul, Frank LW Takken

## Abstract

Nucleotide-binding Leucine Rich repeat-type immune receptors (NLRs) are intracellular proteins that sense the presence of pathogen-derived elicitors and subsequently trigger an immune response. NLR proteins have to be strictly regulated as immune responses typically result in death of affected host cells. Regulation mechanisms of NLR activation are well studied, however steps immediately following NLR activation are largely unexplained. Multimerization of NLRs is thought to be involved, although currently no unambiguous paradigm regarding the dynamics of this process exists. Some NLRs form high-molecular weight complexes before activation, others exclusively after activation, or, like Rx1, none could be detected. We investigated NLR complex formation in transgenic *N. benthamiana* stably expressing the potato Rx1 protein from its native promoter. Activation of the Rx1-resistance response was synchronized by dexamethasone-controlled expression of its elicitor; the Potato Virus X Coat Protein. Rx1 self-associates upon activation: Rx1 homomers are absent before dexamethasone application, a complex could be detected 1 hour after application, but surprisingly is again absent after 2 hours or later. These results show that self-association of NLR proteins upon activation can be transient, explaining the difficulties of detecting them during the normal, non-synchronized infection process as this typically involves few affected cells at any time.

## Introduction

Plants defend themselves against pathogens with a multilayered immune system (Cook et al., 2015; Dodds and Rathjen, 2010). A first line of defense is activated by cell-membrane localized immune receptors that sense extracellularly localized pathogen-derived molecules. The second line of defense is mounted by intracellular immune receptors recognizing molecules that pathogens insert into host cells. A major class of these intracellular receptors is the NLR (Nucleotide-binding Leucine-rich Repeat)-type protein (Li et al., 2015). Activation of NLR receptors upon pathogen perception typically triggers a multi-faceted immune response that includes production of reactive-oxygen species (ROS), various antimicrobial metabolites and pathogenesis related proteins. Furthermore, callose and lignin depositions result in cell wall fortification, and often a Hypersensitive Response (HR) is triggered resulting in death of the infected cells (Cui et al., 2015; Knip et al., 2019).

NLR receptors, and the processes triggered by activated NLRs, have been studied extensively [Reviewed in:(Adachi et al., 2019; Cui et al., 2015; Li et al., 2015; Zhang et al., 2017)]. Based on their N-terminal domains NLRs are generally divided into two major classes, namely, Toll/interleukin-1 receptor/resistance Nucleotide-binding Leucine-rich repeat proteins (TNLs) and Coiled-coil receptor Nucleotide-binding Leucine-rich repeat proteins (CNLs) (Li et al., 2015). The intra- and intermolecular interactions of these proteins that keep them in an inactive, but activatable, state have been studied intensively (Chen et al., 2016; Lukasik-Shreepaathy et al., 2012; Rairdan et al., 2008; Rairdan and Moffett, 2006; Sukarta et al., 2016). Also, other regulatory mechanisms to control NLR activity for instance regulated turnover, localization and stability, have been studied (Borrelli et al., 2018). However, a molecular mechanism to link NLR activation to activation of immune signaling is still lacking. A complicating factor to study NLRs proteins in infected plant tissues is the rapid cell death triggered upon NLR activation (Knip et al., 2019). Furthermore, NLR activation is dependent on pathogen perception, which is rarely, if ever, synchronous across tissues. Infected tissues will contain non-, early- and late-responding cells of which some might have already undergone cell death. Despite the difficulty of studying NLRs *in planta*, they have been found to bind chaperones and to homo/heteromerize. Homomerization of NLRs, or at least of their N-terminal domain, has been reported for NLRs of both TNL and CNL classes (Griebel et al., 2014). Homomerization is essential to trigger cell death for the TNL L6 from flax and the CNL MLA10 from barley (Bernoux et al., 2011; Maekawa et al., 2011). However, NLRs in a pre-activation state have also been reported to homomerize, like the CNLs Prf from tomato and RPS5 from Arabidopsis, raising questions about the dynamics of NLR complex-formation with regards to their activation (Ade et al., 2007; Gutierrez et al., 2010; Ntoukakis et al., 2014). Methods like yeast-two hybrid or co-immunoprecipitation can reveal interactions, but do not provide quantitative insights in the number of NLRs engaged in a complex. Excitingly, the recent elucidation of the first crystal structure of a plant NLR protein in the pre- and post-activation state shows that the CNL ZAR1 forms an ADP-bound monomer before elicitor perception. Upon perception of the PBL2 elicitor the ZAR1-RKS1 complex forms an ATP-bound pentameric ring-like structure, which is proposed to reflect the activated state of the NLR (Wang et al., 2019). Therefore, although some NLRs self-associate in a pre-activation state, higher-order complex formation has been proposed to be a generic mechanism for NLRs to trigger immunity. However, so far only a few NLRs have been reported to form such higher-order complexes *in planta* upon elicitor perception (Bernoux et al., 2011; Casey et al., 2016; Cesari et al., 2016; Maekawa et al., 2011; Mestre and Baulcombe, 2006; Schreiber et al., 2016; Tran et al., 2017). One explanation is that multimerization is not a generic feature involved in NLR activation. An alternative explanation is that technical challenges prevent the detection of such complexes due to e.g. low accumulation levels of the NLR, cell death triggered upon NLR activation or the heterogenous nature of the immune response.

To distinguish between these two possibilities we made use of the CESSNA (Controlled Expression of effectors for Synchronized and Systemic NLR Activation) platform to co-express the potato CNL Rx1 with its elicitor, the Coat Protein (CP) from Potato Virus X (PVX). While Rx1 expression was driven by its endogenous promoter, the expression of CP was controlled by a dexamethasone (DEX)-inducible promoter. Rx1 was chosen because, despite many efforts, self-association upon elicitor perception has not been detected (Slootweg et al., 2018). Here we show that Rx1 forms higher order complexes one hour after inducing *CP* expression by DEX application, but these complexes dissociate again at later time points. These findings imply that NLR-multimerization upon activation might be a generic process, but that oligomerization could be easily missed due to its transient nature. Our results stress the importance of a synchronized system for NLR activation to study NLR multimerization and identify interacting partners.

## Results

To investigate Rx1 self-interaction, co-immunoprecipitation (CoIP) experiments were performed. Plants with *pRx1::Rx1:4HA / DEX::CP106* constructs (referred to as *Rx1D106*), were transiently transformed with a *35SLS::Rx1:GFP* construct by Agroinfiltration of the first fully expanded leaves (Ma et al., 2012; Slootweg et al., 2010). Two days after transient transformation the infiltrated leaves were brushed with DEX to induce CP106 production. Soluble proteins were extracted and immunoprecipitation was done using Chromobeads binding the GFP-tag of the chimeric Rx1:GFP protein. Following SDS-polyacrylamide gel electrophoresis (SDS-PAGE), and immunoblotting using an HA-antibody Rx1:4xHA was readily detected at all timepoints in the input material. Notably, Rx1 was not only found at its expected size (~100kDa for Rx1:4xHA), but also a specific band of an apparent molecular mass well above 180kDa was observed in the Rx1:4xHA input samples. Probing the input material with a GFP-antibody to detect Rx1:GFP did not reveal a signal, indicating that the amount of protein is below the detection level of the antibody, as after immunoprecipitation a specific band was observed on immunoblot (Figure 1). The presence of this band, likely corresponding to Rx1:GFP, shows that immune precipitation was successful. Like with the HA antibody, a second band cross-linking with the antibody was observed with a size >180kDa (Figure 1, blue arrows). When probing the GFP-immunoprecipitants using the HA-antibody a band was only observed in the extract harvested one hour after DEX application, but not in the extract harvested at the 2h timepoint or at T=0. This result indicates that Rx1 forms a homomeric complex after DEX application and CP production, but that at later timepoints this complex dissociates (Figure 1).

**Figure 1.**
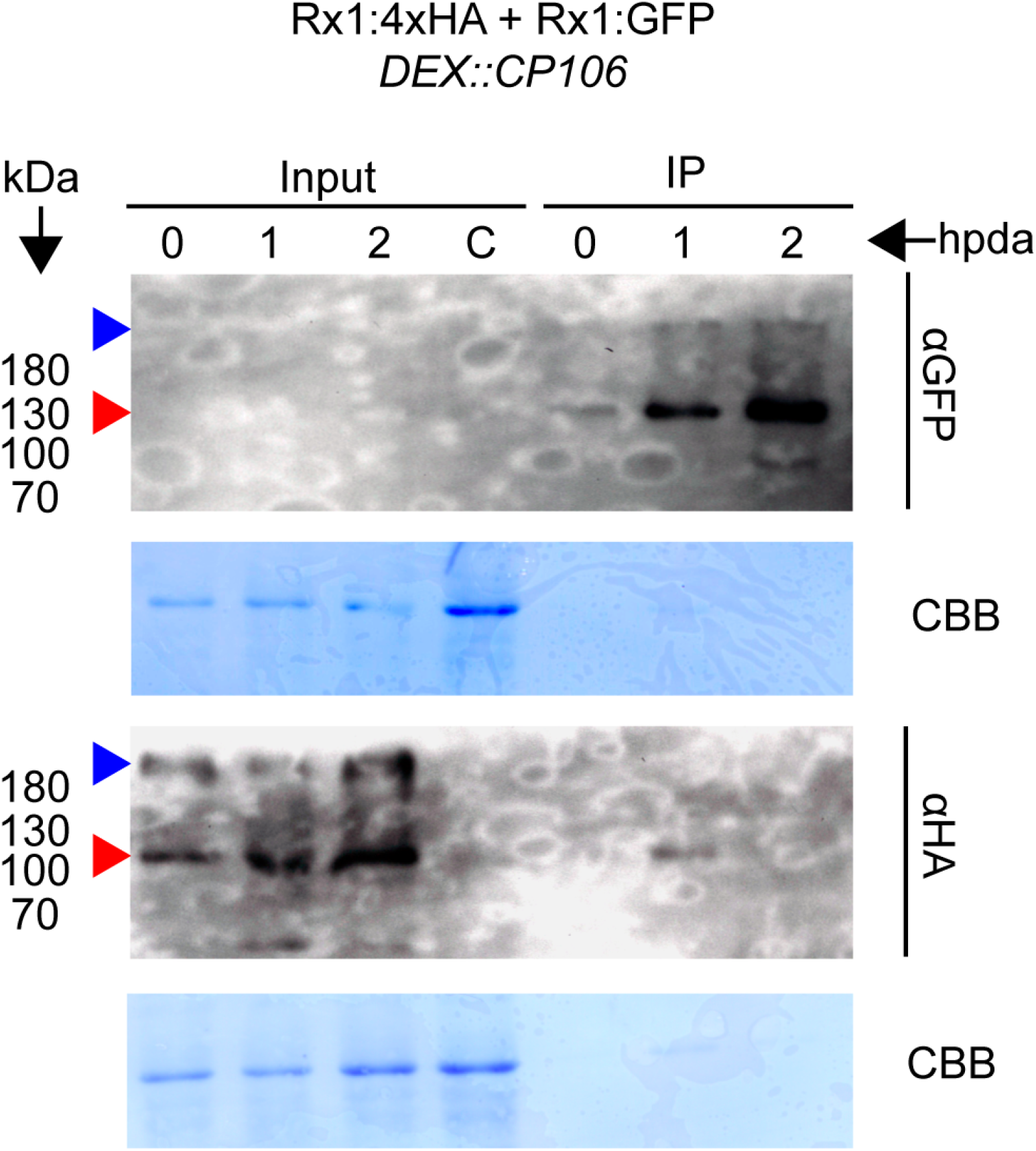
CoIP of Rx1:4HA and Rx1:GFP after induction of CP production. On the top Rx1:GFP is visualized on an SDS page gel using GFP antibody, prior (input) and after IP. Below Rx1:4xHA is detected using HA-antibody, prior to CoIP (input) and after CoIP. Coomassie loading controls are displayed below the SDS PAGE panes. Red arrows indicate the height at which monomeric Rx1 fusion proteins are migrate, blue arrows indicate a band of approximately twice the size of monomeric Rx1.

To investigate the formation and size of the putative Rx1 complexes, native conditions for protein extraction were used and Blue-native polyacrylamide gel electrophoresis was performed (BN-PAGE). Unlike SDS-PAGE, in which complexes are dissociated, BN-PAGE allows identification of high molecular weight complexes. *Rx1D106* plants were treated with DEX to induce expression of CP106. Subsequently, tissue was harvested at 0, 1 and 2 h post DEX induction. As control, a DEX::CP106 plant was used that carries the DEX inducible CP construct, but not Rx1:4xHA, to assess specificity of the HA-antibody. To monitor whether the DEX::CP106 construct affects migration of the Rx1-HA protein, a transgenic line was also used that expresses Rx1::4xHA, but does not contain the DEX::CP106 construct. The samples were divided into two and used to prepare protein extracts for separation on either BN-PAGE or SDS-PAGE. Separation of the protein extracts of the *Rx1D106* plants at the three indicated timepoints using native gels revealed that the bulk of Rx1:4xHA migrates at an apparent mass between 146 and 242kDA. Besides, a smear is visible at higher molecular mass, which implies the presence of a larger protein complex (Figure 2A). This smear is not visible, or to a much lesser extent, in the Rx1:4xHA sample indicating that its presence is triggered by the presence of the DEX:CP106 construct (See also, Figure S1). The absence of a signal in the DEX:CP106 and wt plants shows that the band cross-reacting with the HA antibody is a genuine Rx1-HA signal. Taken together these data suggest that Rx1 forms a higher molecular weight complex in the presence of the CP106 construct.

**Figure 2.**
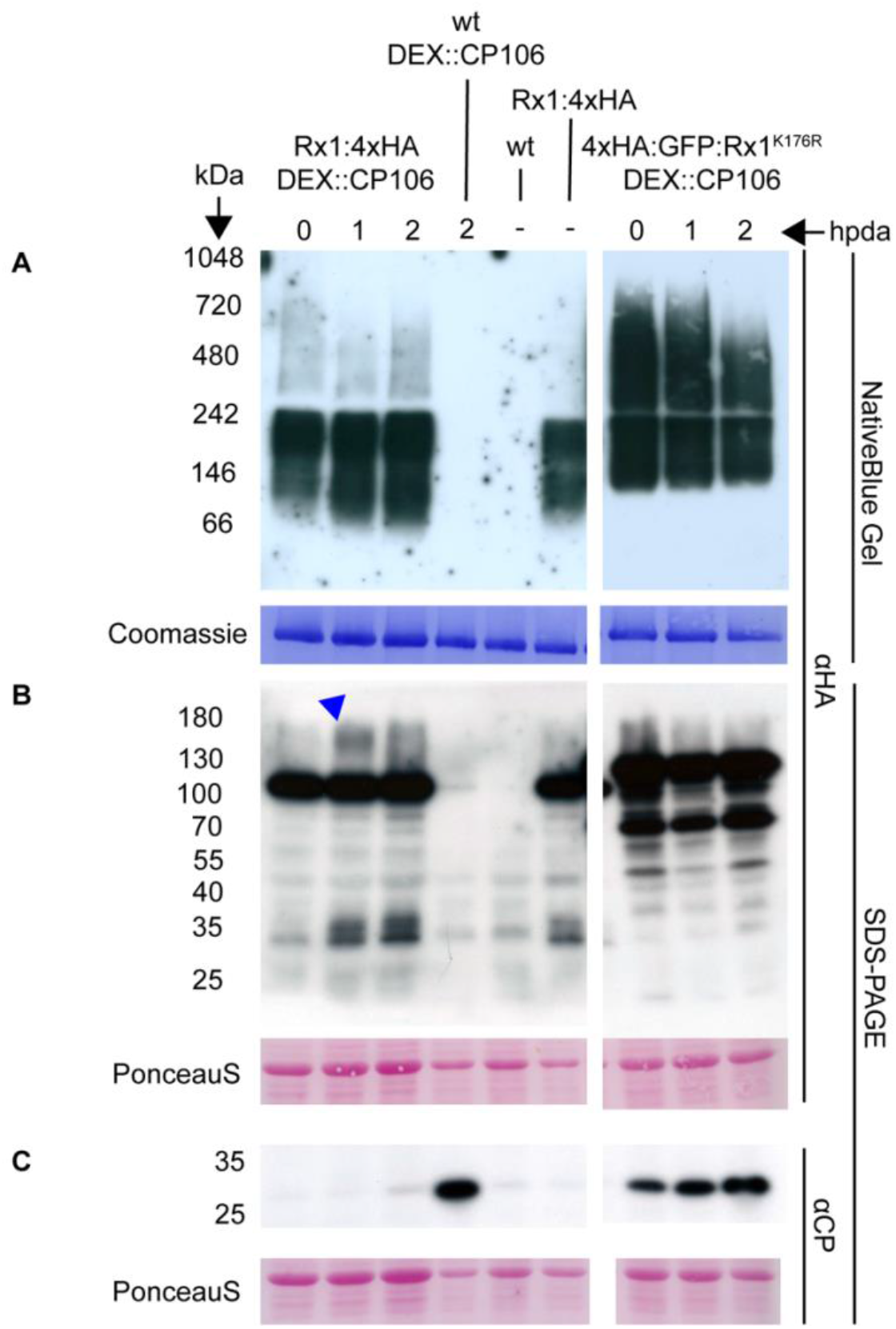
Native Blue- and SDS-PAGE analysis of Rx1 after DEX-triggered induction of CP production. (A) Native blue PAGE probed with an HA-antibody to detect native HA tagged Rx1 protein (upper panel), and Coomassie Brilliant Blue stain of the same blot to show total protein abundance (lower panel). (B) SDS PAGE using an HA-antibody to detect wt and mutant Rx1 protein (upper panel) and a Ponceau S stain of the same membrane serving as loading control (lower panel). Band in sample *Rx1D106* T=1 at ~180kDa is marked with a blue arrow (C) SDS PAGE probed with an anti-PVX antibody to detect CP accumulation (upper panel) and a Ponceau S stain of the membrane serving as loading control (lower panel). As protein abundance varied between the *Rx1D106* and HA:GFP:Rx1_K176R_ samples, Figure 2A is a collage of 2 films made from the same immunoblot. Pictures of the original films can be found in Figure S1.

To test whether formation of this higher molecular weight complex requires a functional Rx1 protein the Rx1_K176R_ mutant was used. This variant carries a mutation in the conserved lysine in the p-loop, which is required for nucleotide binding in related NLR proteins (Tameling et al., 2006). The Rx1_K176R_ mutant is non-responsive to CP and unable to trigger immune responses (Knip et al., 2019; Moffett et al., 2002). Surprisingly, 4xHA:GFP:Rx1_K176R_ was found to accumulate to higher levels than wild-type Rx1:4xHA (Figure 2A), but like wt Rx1 a similar smear of higher molecular weight products was observed.

Analyzing the same samples on SDS-PAGE reveals an apparent size of just over 100kDa for Rx1:4xHA, and of around 130kDA for 4xHA:GFP:Rx1_K176R_ consistent with the predicted sizes of the proteins of 111,5 and 135,5kDa kDa. Moreover, some bands of lower molecular weight are observed in Rx1 and Rx1_K176R_ containing samples that likely represent degradation products (Figure 2B). Whereas the intensity of the breakdown product did not change for the mutant the intensity of breakdown products with a Mw of around 35kDA increased for wt Rx1, indicating a specific response for the latter. In addition, only in the *Rx1D106 (*Rx1:4xHA DEX::CP106) sample an SDS-resistant band of an apparent size of 130 and 180kDa is observed one hour after DEX application that is not present at other time points nor in the other Rx1 or Rx1_K176R_ containing samples (Figure 2B; blue arrow). These observations indicate that complex formation and protein degradation are linked to Rx1 activation.

As Rx1 activation relies on the presence of CP, the protein extracts described above were separated using SDS-PAGE and used for immunoblotting, to monitor the induction of CP upon DEX treatment. Probing the blots with a CP-specific antibody revealed a strong signal in the DEX:CP106 lines and no signal in wt plants confirming the specificity of the antibody (Figure1C). In the presence of 4xHA:GFP:Rx1_K176R_, induction of CP accumulation is observed upon DEX treatment. However, the presence of a signal at T=0 indicates that the protein is also expressed, albeit to lower levels, without DEX, indicating leakiness of the promotor. When DEX is applied to the RX1::HA/DEX::CP106 plants, CP is found to accumulate to very low levels at early time points and become only clearly detectable two hours post DEX application. (Figure 2C). To conclude, CP production is induced upon DEX application, but its accumulation is suppressed in wt Rx1 plants.

## Discussion

Using our DEX-inducible CESSNA platform (Knip et al., 2019) we observed transient Rx1 self-association after induction of CP expression. The use of the CESSNA system was key to reveal homomerization as the interaction was only detectable 1 hdpa and would have been overlooked without a synchronized induction. In addition, only a small fraction of Rx1:4xHA was pulled-down following CoIP, stressing the importance of a spatiotemporal system in which all cells respond simultaneously, as otherwise the low amount of post-activation NLR complexes in the few responding cells would easily prevent their detection (Figure 1). Besides homomeric interactions upon CP perception the Rx1 protein also forms heteromeric interactions in the absence of the Coat Protein. In both anti-HA probed input material and anti-GFP probed IP material an additional band of >180kDa was visible on the immune blots. Since CoIP only reveals a stable interaction of Rx1:4xHA and Rx1:GFP at 1 hpda (Figure 1), this higher molecular weight complex is likely a heteromer, possibly containing (co)chaperones like SGT1 (suppressor of the G2 allele of Skp1), HSP90 (Heat Shock Protein of 90kDa), PP5 (Protein Phosphatase 5) or Hsp20 (Heat Shock Protein of 20kDa) (Bieri et al., 2004; Boter et al., 2007; de la Fuente van Bentem et al., 2005; Van Ooijen et al., 2010). Rx1 is dependent on the SGT1-RAR1-HSP90 complex for its activity, although a direct interaction has not been reported.

Blue Native gels suggest that Rx1 engages in various protein complexes as the protein is detected in a smear over a wide molecular weight range. The specific band of a Mw between 130 and 180kDa is of interest, as it is only detected at 1 hpda implying that its formation is dependent on the activation state of Rx1 (Figure 2B; blue arrow). Given the estimated size it is unlikely to be a dimer of Rx (estimated size 223 kDa), but since the resolution of SDS-PAGE above 100kDa is not very high we cannot fully exclude this possibility. Besides the aforementioned chaperones, NbGlk1 (Golden-like Kinase 1) and RANGAP2 (RAN GTPase Activating Protein 2) have been reported to physically interact with Rx1 (Sacco et al., 2007; Tameling et al., 2010; Townsend et al., 2018). The size of the band could signify Rx1:4xHA bound to RANGAP2 (~171 kDA), or to Glk1 (~158kDa), which would imply that Rx1 binds these proteins in an activation-dependent manner forming a SDS-tolerant complex. Notably, similar bands where not observed in the input samples of the CoIP experiment, which uses a different extraction buffer, suggesting that a component in this buffer disrupts the complex. Epitope-tagged RANGAP2 or GLK1 protein and/or specific antibodies are required to test the hypothesis on whether these specific proteins are engaged in a transient Rx1 complex.

The Rx1_K176R_ protein accumulates to higher levels than the wt Rx1 protein. A possible explanation could be that the miRNAs from the mir482-family and mir6024 are unable to suppress the mRNA accumulation levels of the Rx1_K176R_ variant. These miRNAs target the conserved sequence of the p-loop that contains the K176R mutation (Li et al., 2012; Seo et al., 2018; Shivaprasad et al., 2012; Zhang et al., 2016). In addition to an increase in abundance, the high molecular weight (>242 kDa) smear visible on BN-PAGE was also much more pronounced (Figure 2A), as are the higher number of bands below 100kDa in SDS-PAGE (Figure 2B). The mutant is predicted to be compromised in nucleotide binding, which means that it is likely to adopt a more “open” conformation, as the nucleotide acts as an organizing center in CNLs (Takken and Tameling, 2009; Wang et al., 2019). The inactive open conformation might be more prone to aggregation - explaining the high molecular weight complexes - and to degradation resulting in the higher abundance of the lower molecular weight bands observed (Figure 2AB).

We find that CP accumulation is strongly induced upon DEX treatment in both wild-type and Rx1_K176R_ *N. benthamiana*, but surprisingly not in Rx1 plants (Figure 2C). Although CP accumulation was modest in the Rx1 lines the protein is still expressed to sufficiently high levels to trigger Rx1 activation as *Rx1D106* leaves collapse about four hours after DEX application (Knip et al., 2019). The increase in low molecular weight bands that cross-react with the HA antibody in the *Rx1D106* line upon DEX treatment is another indication that the CP is expressed and changes the Rx1 conformation making it more prone to degradation. The mechanism preventing the accumulation of the CP in Rx1 lines is unknown, but it is tempting to speculate that translational inhibition induced by the activated Rx1 protein is involved. Translational inhibition of CP accumulation has been reported before upon activation of the tobacco N protein by its cognate Tobacco mosaic virus derived elicitor (Bhattacharjee et al., 2009).

The identification of the transient nature of the formation of a Rx1 multimer upon elicitor perception is of relevance for the study of NLR proteins. These dynamics could explain why this Rx1 complex has not been observed before (Slootweg et al., 2018). Whether the formation of other NLR proteins’ complexes is as dynamic as for Rx1 is unknown, but if so, then an inducible system to systemically activate immune receptor is required to capture its presence. Although a post-activation NLR oligomeric complex produced in an heterologous system is sufficiently stable to allow its structural elucidation (Wang et al., 2019), the lifetime of such an NLR complex in intact leaf tissues could be much shorter, possibly due to the induction of the immune responses triggered by those proteins.

## Methods

### Plant line generation, transient transformation and growth condition

Stable transgenic *N. benthamiana* lines were generated using a method adapted from Sparkes *et al*. (Sparkes et al., 2006). *Rx1:4xHA N. benthamiana* plants were transformed with pTA7002 Gateway DEX::CP106 (Knip et al., 2019; Lu et al., 2003), and wild-type N. benthamiana was transformed with pBIN+ *35SLS::4HA:GFP:Rx1_K176R_* construct (Slootweg et al., 2010). After Agroinfiltration leaf sectors were incubated on MS (Murashige and Skoog)- plates containing 40μg/ml Hygromycin (for DEX::CP106 containing plants) or 100μg/ml kanamycin (for *35SLS::4HA:GFP:Rx1_K176R_*). Emerging antibiotic resistant plantlets were transferred to soil to allow flowering and seed set. Progeny of these plants (T_2_) were checked by PCR (Kanamycin (NPTII) and Hygromycin (HPT)-gene specific primers, see Table S1) for presence of the *DEX::CP106/35SLS::4HA:GFP:Rx1_K176R_*. Segregation of the transgenes was determined by germinating the seeds on MS-media containing 40μg/ml Hygromycin or 100μg/ml Kanamycin. Zygosity of the transgenes was determined using quantitative PCR to select homozygous plants as described (Glowacka et al., 2016) (See Table S1 for NPTII and HPT primers and for a internal reference gene (NRG1)). Plants carrying *pRx1::Rx1:4HA / DEX::CP106* constructs are referred to as *Rx1D106* (lab stock reference #FP1807). Wild-type *N. benthamiana* plant-line containing the pBIN+ *35SLS::4HA:GFP:Rx1_K176R_* construct as referred to as *Rx1_K176R_* (lab stock reference #FP1888). *N. benthamiana* were grown in long-day conditions in a climate chamber (22°C, 70% humidity, 11h/13h light/dark). Agrobacterium mediated transient transformation was performed on the youngest fully expanded leaves of 4-5 week old plants as described (Ma et al., 2012).

### Plasmid construction

The pTA7002 Gateway DEX::CP106 construct has been described (Knip et al., 2019). A Gateway^™^-compatible version of pBIN+ adding a GFP tag, called pBINLR+ GFP (pFP1806), was generated by cloning the expression module from pK7FWG2 (Karimi et al., 2002) into the SmaI site of pBIN+ (Table S1) (van Engelen et al., 1995). pBINLR+ 35SLS::Rx1:GFP was generated by PCR amplifying the Rx1 CDS (FP7899 and FP7902), cloning it into pDONR207, which was used to recombine the Rx1 CDS into pBINLR+ GFP (pSDM6064) (Table S1). Importantly, primer FP7899 adds a secondary ATG-site just before the transcription start site of Rx1, giving rise to a “leaky-scan” (LS) construct as described (Slootweg et al., 2010). This ensures constitutive expression of Rx1 at low levels. The pBIN+ 35SLS::Rx1:GFP:HA, 35SLS::Rx1_D245E_: GFP:HA and 35SLS::4xHA:GFP:Rx1_K176R_ constructs are described (Slootweg et al., 2010).

### Native Protein isolation and Native Blue PAGE

Proteins were isolated from six leaf discs (5mm Ø World Precision Instruments (WPI)), by placing the discs in a 2ml Eppendorf tube and grinding them using 3mm Ø metal beads and a Qiagen TissueLyzer II at 60hz for 30 sec. in 100μl GTEN-buffer (10% Glycerol, 25mM TRIS pH 8, 0.5mM EDTA, 150 mM NaCl,1 x protease inhibitors (Roche), 5mM DTT, 1% NP40) on ice. Samples were then spun down at 4_O_C and 13000g for 5 minutes. Per sample 15μl of the supernatant was used and 6μl 4x Sample Buffer, 1 μl Coomassie Brilliant Blue G-250 and 2 μl water were added according to the manufacturer’s instructions (NativePageTM Bis-Tris Mini gel electrophoresis protocol, Thermo-Fisher Scientific). Samples were loaded on a NativePAGE^™^ 3-12% Bis-Tris Protein Gel (Thermo-Fisher Scientific) alongside a NativeMark^™^ Unstained Protein Standard (Thermo-Fisher Scientific). Electrophoresis was performed at 150V in a Mini Gel Tank (Thermo-Fisher Scientific). After electrophoresis proteins were transferred to a Polyvinylidene fluoride (PVDF; Immobilon-P^™^, Millipore®) membrane using semi-dry blotting (Amersham biosciences TE 77 ECL Semi-Dry Transfer Unit) in combination with the NuPageTM transfer buffer (Thermo-Fisher Scientific). After protein transfer the PVDF membrane was incubated in 8% Acetic Acid for 15 minutes to fix the proteins and subsequently the blots where rinsed with distilled water before probing with epitope tag-specific antibodies.

### Co-immunoprecipitation

For co-immunoprecipitation (coIP) analysis, four to five-week old *Rx1D106 N. benthamiana* plants were transiently transformed with pBINLR+ 35SLS::Rx1:GFP. An *A. tumefaciens* GV3101 over-night culture (OD_600_ 0.7-1.5) containing pBINLR+ 35SLS::Rx1:GFP (pSDM6152) was harvested and the cells resuspended in infiltration buffer and diluted to OD_600_ 0.05 (Ma et al., 2012). This suspension was infiltrated in the two upper fully expanded leaves. Two days after infiltration, CP106 production was induced by brushing DEX on the leaves with a paintbrush (20 μM DEX, 0.01% Silwet in 15 mL miliQ). Leaves were harvested at 0, 1 and 2 hours after by brushing DEX and snap frozen in liquid nitrogen. Un-infiltrated leaves and leaves untreated with DEX were used as controls for infiltration and DEX-treatment. Frozen tissue was ground until a fine powder using a mortar and pestle using liquid nitrogen. To remove phenolic compounds 1% Polyvinylpolypyrrolidone was added whilst grinding. A GTEN-based buffer (10% Glycerol, 25mM TRIS pH 8, 0.5mM EDTA, 150 mM NaCl, 1 x protease inhibitors (Roche), 5mM DTT, 1% NP40) was used for protein extraction. Extraction buffer was added in a 1:2 g/mL ratio to ground leaf material. Samples were thawed on ice and sonicated three times 10 seconds, with 10 seconds intervals in between. Subsequently, samples were transferred to 2mL Eppendorf tubes and centrifuged at 4°C at 13000 rpm for 15 minutes. As control, 40 μL supernatant was taken and stored at −20°C; hereafter referred to as ‘input’. Subsequently, 1.5-2 mL filtered (Millex GP filter unit, 0.22μm, PES membrane, MilliporeSigma^™^) supernatant was mixed with GFP-Trap® magnetic beads (Chromotek®) (15 μL beads per sample, washed three times with wash buffer) and incubated on a roller mixer for 1 hour at 4°C. GTEN-based wash buffer consisted of 10% Glycerol, 25mM Tris pH 8, 0.5mM EDTA, 150 mM NaCl, 1 x protease inhibitors (Roche®). Beads were pelleted using a magnetic separator, washed three times and resuspended in 30μl wash buffer. 10 μL 5X Laemmli buffer (375 mM Tris-HCl pH 6.8, 6% SDS, 20% v/v glycerol, 0.03% bromophenol blue and 100 mM DTT) was added to input and coimmunoprecipitated samples, heated at 95°C for 5 minutes and analyzed by SDS-PAGE and immuno blotting.

### SDS PAGE and immunodetection

For the detection of Rx1:GFP fusion proteins, proteins were isolated from six leaf discs (5mm Ø). The discs were placed in a 2ml Eppendorf tube and homogenizing them using 3mm Ø metal beads and a Qiagen TissueLyzer II at 60hz for 30 sec. in 100μl extraction buffer (Tris-HCl 50 mM pH 6.8, SDS 2%, DTT 2 mM and 1× protease inhibitors (Roche) and centrifuged for 5 min at 13,000g at 4_O_C. The supernatant was mixed 5:1 with 5× Laemmli buffer (375 mM Tris-HCl pH 6.8, 6% SDS, 20% v/v glycerol, 0.03% bromophenol blue and 100 mM DTT) and heated at 95°C for 5 minutes. Total proteins were separated on 10% SDS-PAGE and (semi-dry) blotted onto Polyvinylidene fluoride (Immobilon-P^™^, Millipore®) membranes.

For the detection of Coat Protein in higher order complexes, the extraction method used for co-immunoprecipitation was used. After extraction, samples were mixed with Laemmli buffer, boiled for 2 minutes and separated on 10% SDS-PAGE and (semi-dry) blotted onto Polyvinylidene fluoride (Immobilon-P^™^, Millipore®) membranes.

PVX-CP was detected using a rabbit PVX-specific polyclonal antibody (diluted 1:3,000) (ref. 110411, Bioreba, Reinach, Switzerland), followed by incubation with horseradish peroxidase (HRP)-conjugated goat anti-rabbit IgG secondary antibody (diluted 1:10,000) (ref. 31460, Pierce^™^). GFP was detected using a rat 3H9 GFP-specific polyclonal antibody (diluted 1:1000)(ref 3H9-100ab, Chromotek, Germany, followed by incubation with horseradish peroxidase (HRP)-conjugated goat anti-rat IgG secondary antibody (diluted 1:10,000)(ref. 31470, Pierce^™^). Secondary immunoglobulins were visualized using home-made ECL solution containing 2.5 mM luminol, 0.4 mM p-coumaric acid, 100mM Tris-HCl pH 8.5 and 0.018% H_2_O_2_. Incubation of both primary and secondary antibodies was done in Tris-buffered saline with 0.05% Tween-20 (TBST) followed by three rinses of 10 minutes in TBST. Equal protein loading was confirmed for the samples by Ponceau-S staining of the membranes.

## Author Contributions

MK and EL performed the experiments. MK performed cloning and generated plant lines. MK performed the Native-PAGE experiments and EL performed the CoIPs. MK and FLWT wrote the manuscript.

## Acknowledgements

We would like to thank Dr. Erik Slootweg for providing Rx1 and mutant derivative constructs, and Tieme Helderman for his help with co-immunoprecipitations. We thank Machiel Beijaert for help with generating transgenic lines. We are also grateful for critical reading and feedback on the manuscript by Martijn Rep. MK, EL and FLWT received funding from the NWO-Earth and Life Sciences funded VICI project No. 865.14.003. The authors declare to have no conflicts of interest.

---

**Supplemental Figure S1.**
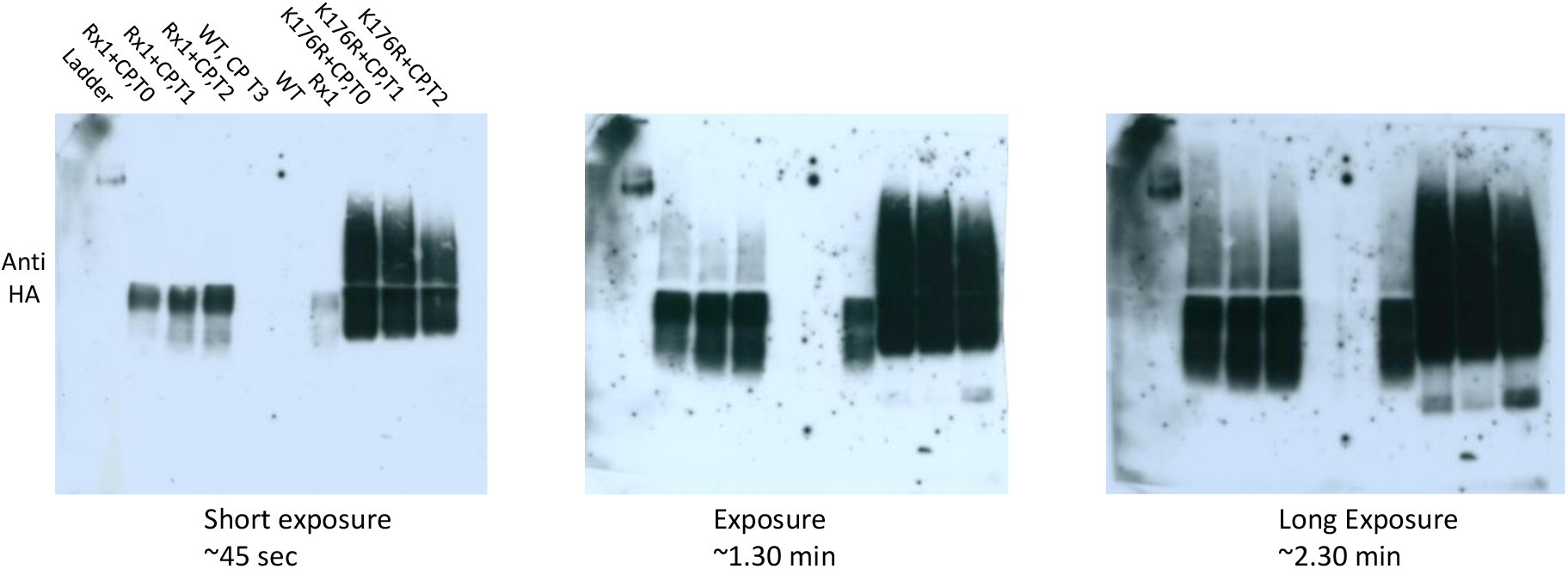
Native Blue Immunoblots (Depicted in Figure 2) visualized with different exoosure times

**Supplemental Table T1.**
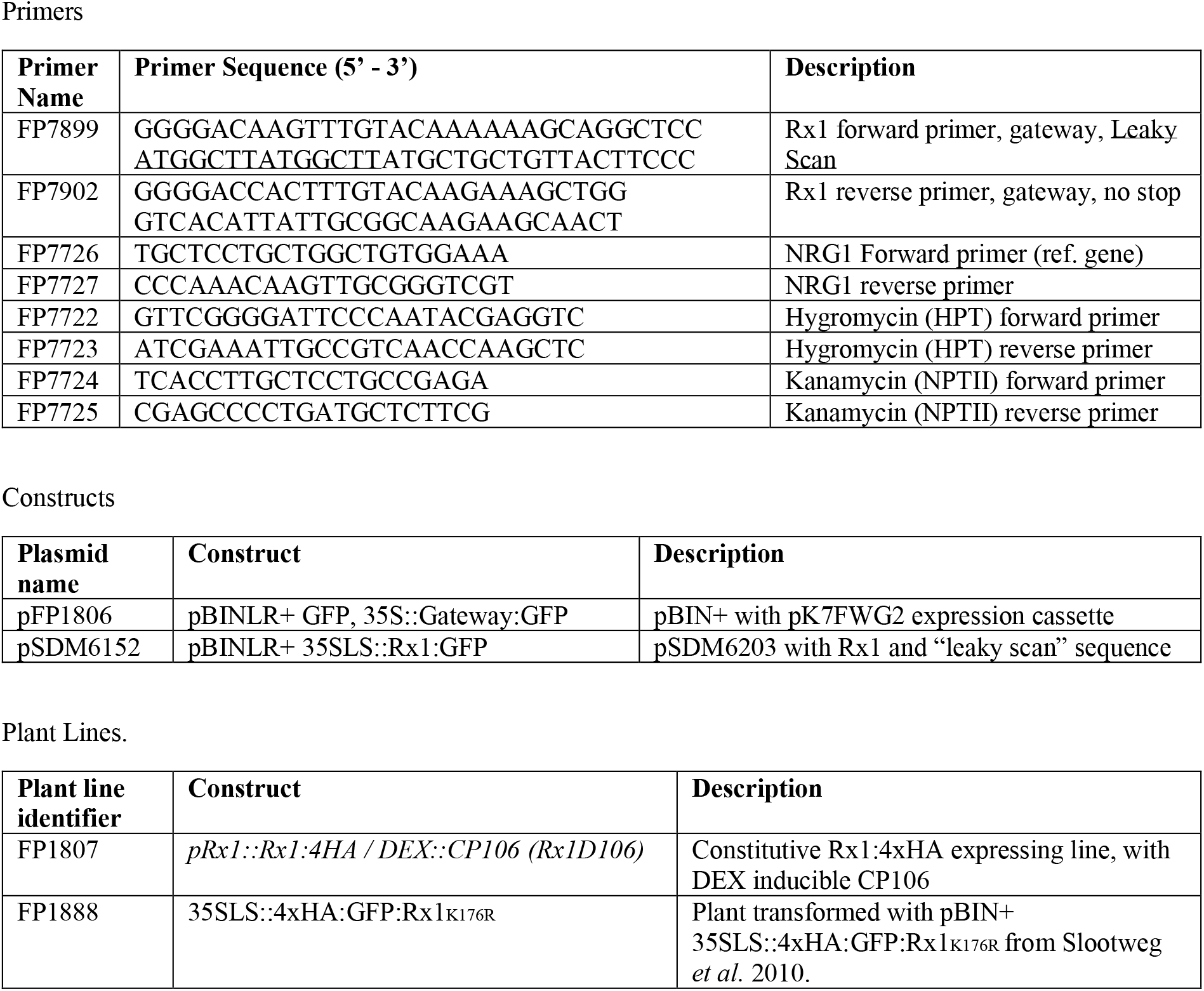
This table contains all the primers used for cloning and (quantitative) PCR (See:”Primers”). It contains the plasmids used the manuscript (See:”Constructs”) and the transgenic homozygous *N. Benthamiana* plant lines (See: “Plant Lines”).

## References

Adachi, H., Derevnina, L., and Kamoun, S. (2019). NLR singletons, pairs, and networks: evolution, assembly, and regulation of the intracellular immunoreceptor circuitry of plants. Curr Opin Plant Biol 50:121–131.

Ade, J., DeYoung, B.J., Golstein, C., and Innes, R.W. (2007). Indirect activation of a plant nucleotide binding site-leucine-rich repeat protein by a bacterial protease. Proc Natl Acad Sci U S A 104:2531–2536.

Bernoux, M., Ve, T., Williams, S., Warren, C., Hatters, D., Valkov, E., Zhang, X., Ellis, J.G., Kobe, B., and Dodds, P.N. (2011). Structural and functional analysis of a plant resistance protein TIR domain reveals interfaces for self-association, signaling, and autoregulation. Cell Host Microbe 9:200–211.

Bhattacharjee, S., Zamora, A., Azhar, M.T., Sacco, M.A., Lambert, L.H., and Moffett, P. (2009). Virus resistance induced by NB-LRR proteins involves Argonaute4-dependent translational control. Plant J 58:940–951.

Bieri, S., Mauch, S., Shen, Q.H., Peart, J., Devoto, A., Casais, C., Ceron, F., Schulze, S., Steinbiss, H.H., Shirasu, K., et al. (2004). RAR1 positively controls steady state levels of barley MLA resistance proteins and enables sufficient MLA6 accumulation for effective resistance. Plant Cell 16:3480–3495.

Borrelli, G.M., Mazzucotelli, E., Marone, D., Crosatti, C., Michelotti, V., Vale, G., and Mastrangelo, A.M. (2018). Regulation and Evolution of NLR Genes: A Close Interconnection for Plant Immunity. Int J Mol Sci 19.

Boter, M., Amigues, B., Peart, J., Breuer, C., Kadota, Y., Casais, C., Moore, G., Kleanthous, C., Ochsenbein, F., Shirasu, K., et al. (2007). Structural and functional analysis of SGT1 reveals that its interaction with HSP90 is required for the accumulation of Rx, an R protein involved in plant immunity. Plant Cell 19:3791–3804.

Casey, L.W., Lavrencic, P., Bentham, A.R., Cesari, S., Ericsson, D.J., Croll, T., Turk, D., Anderson, P.A., Mark, A.E., Dodds, P.N., et al. (2016). The CC domain structure from the wheat stem rust resistance protein Sr33 challenges paradigms for dimerization in plant NLR proteins. Proc Natl Acad Sci U S A 113:12856–12861.

Cesari, S., Moore, J., Chen, C., Webb, D., Periyannan, S., Mago, R., Bernoux, M., Lagudah, E.S., and Dodds, P.N. (2016). Cytosolic activation of cell death and stem rust resistance by cereal MLA-family CC-NLR proteins. Proc Natl Acad Sci U S A 113:10204–10209.

Chen, X., Zhu, M., Jiang, L., Zhao, W., Li, J., Wu, J., Li, C., Bai, B., Lu, G., Chen, H., et al. (2016). A multilayered regulatory mechanism for the autoinhibition and activation of a plant CC-NB-LRR resistance protein with an extra N-terminal domain. New Phytol 212:161–175.

Cook, D.E., Mesarich, C.H., and Thomma, B.P. (2015). Understanding plant immunity as a surveillance system to detect invasion. Annu Rev Phytopathol 53:541–563.

Cui, H., Tsuda, K., and Parker, J.E. (2015). Effector-triggered immunity: from pathogen perception to robust defense. Annu Rev Plant Biol 66:487–511.

de la Fuente van Bentem, S., Vossen, J.H., de Vries, K.J., van Wees, S., Tameling, W.I., Dekker, H.L., de Koster, C.G., Haring, M.A., Takken, F.L., and Cornelissen, B.J. (2005). Heat shock protein 90 and its co-chaperone protein phosphatase 5 interact with distinct regions of the tomato I-2 disease resistance protein. Plant J 43:284–298.

Dodds, P.N., and Rathjen, J.P. (2010). Plant immunity: towards an integrated view of plant-pathogen interactions. Nat Rev Genet 11:539–548.

Glowacka, K., Kromdijk, J., Leonelli, L., Niyogi, K.K., Clemente, T.E., and Long, S.P. (2016). An evaluation of new and established methods to determine T-DNA copy number and homozygosity in transgenic plants. Plant Cell Environ 39:908–917.

Griebel, T., Maekawa, T., and Parker, J.E. (2014). NOD-like receptor cooperativity in effector-triggered immunity. Trends Immunol 35:562–570.

Gutierrez, J.R., Balmuth, A.L., Ntoukakis, V., Mucyn, T.S., Gimenez-Ibanez, S., Jones, A.M., and Rathjen, J.P. (2010). Prf immune complexes of tomato are oligomeric and contain multiple Pto-like kinases that diversify effector recognition. Plant J 61:507–518.

Karimi, M., Inze, D., and Depicker, A. (2002). GATEWAY vectors for Agrobacterium-mediated plant transformation. Trends Plant Sci 7:193–195.

Knip, M., Richard, M.M.S., Oskam, L., van Engelen, H.T.D., Aalders, T., and Takken, F.L.W. (2019). Activation of immune receptor Rx1 triggers distinct immune responses culminating in cell death after 4 hours. Mol Plant Pathol 20:575–588.

Li, F., Pignatta, D., Bendix, C., Brunkard, J.O., Cohn, M.M., Tung, J., Sun, H., Kumar, P., and Baker, B. (2012). MicroRNA regulation of plant innate immune receptors. Proc Natl Acad Sci U S A 109:1790–1795.

Li, X., Kapos, P., and Zhang, Y. (2015). NLRs in plants. Curr Opin Immunol 32:114–121.

Lu, R., Malcuit, I., Moffett, P., Ruiz, M.T., Peart, J., Wu, A.J., Rathjen, J.P., Bendahmane, A., Day, L., and Baulcombe, D.C. (2003). High throughput virus-induced gene silencing implicates heat shock protein 90 in plant disease resistance. EMBO J 22:5690–5699.

Lukasik-Shreepaathy, E., Slootweg, E., Richter, H., Goverse, A., Cornelissen, B.J., and Takken, F.L. (2012). Dual regulatory roles of the extended N terminus for activation of the tomato MI-1.2 resistance protein. Mol Plant Microbe Interact 25:1045–1057.

Ma, L., Lukasik, E., Gawehns, F., and Takken, F.L. (2012). The use of agroinfiltration for transient expression of plant resistance and fungal effector proteins in Nicotiana benthamiana leaves. Methods Mol Biol 835:61–74.

Maekawa, T., Cheng, W., Spiridon, L.N., Toller, A., Lukasik, E., Saijo, Y., Liu, P., Shen, Q.H., Micluta, M.A., Somssich, I.E., et al. (2011). Coiled-coil domain-dependent homodimerization of intracellular barley immune receptors defines a minimal functional module for triggering cell death. Cell Host Microbe 9:187–199.

Mestre, P., and Baulcombe, D.C. (2006). Elicitor-mediated oligomerization of the tobacco N disease resistance protein. Plant Cell 18:491–501.

Moffett, P., Farnham, G., Peart, J., and Baulcombe, D.C. (2002). Interaction between domains of a plant NBS-LRR protein in disease resistance-related cell death. EMBO J 21:4511–4519.

Ntoukakis, V., Saur, I.M., Conlan, B., and Rathjen, J.P. (2014). The changing of the guard: the Pto/Prf receptor complex of tomato and pathogen recognition. Curr Opin Plant Biol 20:69–74.

Rairdan, G.J., Collier, S.M., Sacco, M.A., Baldwin, T.T., Boettrich, T., and Moffett, P. (2008). The coiled-coil and nucleotide binding domains of the Potato Rx disease resistance protein function in pathogen recognition and signaling. Plant Cell 20:739–751.

Rairdan, G.J., and Moffett, P. (2006). Distinct domains in the ARC region of the potato resistance protein Rx mediate LRR binding and inhibition of activation. Plant Cell 18:2082–2093.

Sacco, M.A., Mansoor, S., and Moffett, P. (2007). A RanGAP protein physically interacts with the NB-LRR protein Rx, and is required for Rx-mediated viral resistance. Plant J 52:82–93.

Schreiber, K.J., Bentham, A., Williams, S.J., Kobe, B., and Staskawicz, B.J. (2016). Multiple Domain Associations within the Arabidopsis Immune Receptor RPP1 Regulate the Activation of Programmed Cell Death. PLoS Pathog 12:e1005769.

Seo, E., Kim, T., Park, J.H., Yeom, S.I., Kim, S., Seo, M.K., Shin, C., and Choi, D. (2018). Genome-wide comparative analysis in Solanaceous species reveals evolution of microRNAs targeting defense genes in Capsicum spp. DNA Res 25:561–575.

Shivaprasad, P.V., Chen, H.M., Patel, K., Bond, D.M., Santos, B.A., and Baulcombe, D.C. (2012). A microRNA superfamily regulates nucleotide binding site-leucine-rich repeats and other mRNAs. Plant Cell 24:859–874.

Slootweg, E., Roosien, J., Spiridon, L.N., Petrescu, A.J., Tameling, W., Joosten, M., Pomp, R., van Schaik, C., Dees, R., Borst, J.W., et al. (2010). Nucleocytoplasmic distribution is required for activation of resistance by the potato NB-LRR receptor Rx1 and is balanced by its functional domains. Plant Cell 22:4195–4215.

Slootweg, E.J., Spiridon, L.N., Martin, E.C., Tameling, W.I.L., Townsend, P.D., Pomp, R., Roosien, J., Drawska, O., Sukarta, O.C.A., Schots, A., et al. (2018). Distinct Roles of Non-Overlapping Surface Regions of the Coiled-Coil Domain in the Potato Immune Receptor Rx1. Plant Physiol 178:1310–1331.

Sparkes, I.A., Runions, J., Kearns, A., and Hawes, C. (2006). Rapid, transient expression of fluorescent fusion proteins in tobacco plants and generation of stably transformed plants. Nat Protoc 1:2019–2025.

Sukarta, O.C.A., Slootweg, E.J., and Goverse, A. (2016). Structure-informed insights for NLR functioning in plant immunity. Semin Cell Dev Biol 56:134–149.

Takken, F.L., and Tameling, W.I. (2009). To nibble at plant resistance proteins. Science 324:744–746.

Tameling, W.I., Nooijen, C., Ludwig, N., Boter, M., Slootweg, E., Goverse, A., Shirasu, K., and Joosten, M.H. (2010). RanGAP2 mediates nucleocytoplasmic partitioning of the NB-LRR immune receptor Rx in the Solanaceae, thereby dictating Rx function. Plant Cell 22:4176–4194.

Tameling, W.I., Vossen, J.H., Albrecht, M., Lengauer, T., Berden, J.A., Haring, M.A., Cornelissen, B.J., and Takken, F.L. (2006). Mutations in the NB-ARC domain of I-2 that impair ATP hydrolysis cause autoactivation. Plant Physiol 140:1233–1245.

Townsend, P.D., Dixon, C.H., Slootweg, E.J., Sukarta, O.C.A., Yang, A.W.H., Hughes, T.R., Sharples, G.J., Palsson, L.O., Takken, F.L.W., Goverse, A., et al. (2018). The intracellular immune receptor Rx1 regulates the DNA-binding activity of a Golden2-like transcription factor. J Biol Chem 293:3218–3233.

Tran, D.T.N., Chung, E.H., Habring-Muller, A., Demar, M., Schwab, R., Dangl, J.L., Weigel, D., and Chae, E. (2017). Activation of a Plant NLR Complex through Heteromeric Association with an Autoimmune Risk Variant of Another NLR. Curr Biol 27:1148–1160.

van Engelen, F.A., Molthoff, J.W., Conner, A.J., Nap, J.P., Pereira, A., and Stiekema, W.J. (1995). pBINPLUS: an improved plant transformation vector based on pBIN19. Transgenic Res 4:288–290.

Van Ooijen, G., Lukasik, E., Van Den Burg, H.A., Vossen, J.H., Cornelissen, B.J., and Takken, F.L. (2010). The small heat shock protein 20 RSI2 interacts with and is required for stability and function of tomato resistance protein I-2. Plant J 63:563–572.

Wang, J., Hu, M., Wang, J., Qi, J., Han, Z., Wang, G., Qi, Y., Wang, H.W., Zhou, J.M., and Chai, J. (2019). Reconstitution and structure of a plant NLR resistosome conferring immunity. Science 364.

Zhang, X., Dodds, P.N., and Bernoux, M. (2017). What Do We Know About NOD-Like Receptors in Plant Immunity? Annu Rev Phytopathol 55:205–229.

Zhang, Y., Xia, R., Kuang, H., and Meyers, B.C. (2016). The Diversification of Plant NBS-LRR Defense Genes Directs the Evolution of MicroRNAs That Target Them. Mol Biol Evol 33:2692–2705.

